# A discontinuous Galerkin model for fluorescence loss in photobleaching of intracellular polyglutamine protein aggregates

**DOI:** 10.1101/303107

**Authors:** Christian V Hansen, Hans J Schroll, Daniel Wüstner

## Abstract

**Background:** Intracellular phase separation and aggregation of proteins with extended poly-glutamine (polyQ) stretches are hallmarks of various age-associated neurodegenerative diseases. Progress in our understanding of such processes heavily relies on quantitative fluorescence imaging of suitably tagged proteins. Fluorescence loss in photobleaching (FLIP) is particularly well-suited to study the dynamics of protein aggregation in cellular models of Chorea Huntington and other polyQ diseases, as FLIP gives access to the full spatio-temporal profile of intensity changes in the cell geometry. In contrast to other methods, also dim aggregates become visible during time evolution of fluorescence loss in cellular compartments. However, methods for computational analysis of FLIP data are sparse, and transport models for estimation of transport and diffusion parameters from experimental FLIP sequences are missing.

**Results:** In this paper, we present a computational method for analysis of FLIP imaging experiments of intracellular polyglutamine protein aggregates also called inclusion bodies (IBs). By this method, we are able to determine the diffusion constant and nuclear membrane permeability coefficients of polyQ proteins as well as the exchange rates between aggregates and the cytoplasm. Our method is based on a reaction-diffusion multi-compartment model defined on a mesh obtained by segmentation of the cell images from the FLIP sequence. The discontinuous Galerkin (DG) method is used for numerical implementation of our model in FEniCS, which greatly reduces the computing time. The method is applied to representative experimental FLIP sequences, and consistent estimates of all transport parameters are obtained.

**Conclusions:** By directly estimating the transport parameters from live-cell image sequences using our new computational FLIP approach surprisingly fast exchange dynamics of mutant Huntingtin between cytoplasm and dim IBs could be revealed. This is likely relevant also for other polyQ diseases. Thus, our method allows for quantifying protein dynamics at different stages of the protein aggregation process in cellular models of neurodegeneration.

## Background

Our understanding of protein transport and aggregation has been revolutionalized by the development of genetically encoded fluorescent protein tags combined with technical innovations in high-resolution live cell fluorescence imaging. In particular, various advanced imaging methods have been used to study aggregation and phase partitioning of proteins in the nucleus and cytosol. Such protein segregation and aggregation is a hallmark of various age-associated neurodegenerative diseases, such as Alzheimer’s disease, Chorea Huntington, Ataxia or Parkinson disease. In several inherited neurodegenerative diseases, like ataxia and Huntington disease, certain proteins bearing a CAG triplet expansion coding for an extended poly-glutamine (polyQ) stretch causes the affected proteins to show the tendency to self-associate and form small and large aggregates, the latter also called inclusion bodies (IBs). Formation of IBs has been associated with disease progression, but it remains unclear, whether such large aggregates are cytoprotective or cytotoxic [1, 2, 3]. In Huntington disease, the polyQ protein is mutated huntingtin (mtHtt) containing more than 30 glutamine repeats typically, while in ataxia, one finds one out of various ataxin proteins mutated containing a polyQ stretch.

The aggregation process in Huntington disease and related polyQ diseases has been studied extensively. Typically, suitable model cells are transfected with fluorescent protein tagged derivatives of the studied polyQ protein, and the aggregation process is studied by a variety of methods including photobleaching techniques like fluorescence recovery after photobleaching (FRAP) and fluoprescence loss in photobleaching (FLIP) [4, 5, 6, 7], number and brightness (N & B) analysis of intensity fluctuations [8], fluorescence complementation assays with split GFP [9], Förster resonance energy transfer (FRET) [6, 4, 10], fluorescence correlation spectroscopy [10], fluorescence lifetime microscopy [4, 11], fluorescence anisotropy imaging [12], stimulated emission depletion (STED) microscopy [13] or single molecule tracking (SMT) [14, 15, 13]. Using such techniques, different aspects of the aggregation process have been revealed. In particular, it has been suggested that diffusive oligomers and small fibrillary aggregates co-exist with IBs, which accumulate after some delay as clearly discernable micron-sized structures [16, 17, 8, 13, 18]. The oligomers or protein fibrils are sometimes difficult to detect, first due to their small size compared to IBs and second due to their low brightness which makes that they are often overshined by the much brighter IBs [8, 15, 13]. However, also the micron-sized IBs formed of green fluorescent protein-tagged mtHtt (GFP-mtHtt) come in strongly varying brightness levels and are eventually preceded by similarly sized but much more dynamic and eventually less bright intermediate structures in the aggregation process [15, 13]. Indeed, protein aggregates detected in cellular models of polyQ diseases are dynamic entities, often recruiting other proteins and thereby sequestering enzymes and signaling proteins which strongly affect the functionality of cells [5, 9, 6, 7]. In detailed FRAP and FLIP studies, both fast- and slow exchanging components have been described for various ataxins and mtHtt with half-times for exchange of tagged protein between cytoplasm and IBs in the range of less than 10-20 sec for various ataxins [19, 20] over 1-2 min for larger IBs of mtHtt6 [4, 20]. This strongly suggests that different populations of inclusions with different physico-chemical properties coexist in affected cells. Supporting that notion, both fibrillary and globular IBs have been detected upon expression of fluorescent protein-tagged mtHtt in the same cells, and this structural heterogeneity was reflected in differing exchange dynamics [4]. An additional level of complexity comes from the complex architecture of the cytoplasm, which generates sub-compartments of varying composition not only via membrane-bound organelles but also in the form of membrane-less liquid phases into which proteins can partition differently [21]. It has been suggested that such variety of physico-chemical phases in the cyto- and nucleoplasm can be a driving force for protein segregation, and in case of mutated polyQ proteins, trigger protein aggregation [22].

Aggregates of polyQ proteins can form in both, the cytoplasm and nucleus, and some polyQ proteins, such as mtHtt or ataxins have been shown to bear nuclear localization and export signals, suggesting active transport across the nuclear membrane [23, 24, 25, 26]. On the other hand for mtHtt, a Ran-GTPase independent transport across the nuclear membrane has been described [27]. How the nucleo-cytoplasmic transport of polyQ proteins is kinetically coupled to their intracellular diffusion and binding to IBs is not known. FLIP is in principle an ideal method to answer this question, as fluorescence loss in different cellular areas can be quantified for repeated localized bleaching far from IBs. However, most studies applying FLIP in this context do not attempt to develop a physical model underlying the observed fluorescence loss kinetics [19, 5, 6]. In a previous study, we presented the first attempt at developing a quantitative FLIP model to estimate exchange rate constants for GFP-mtHtt from FLIP image sequences [7]. We tracked individual IBs and determined exchange rate constants relative to the overall fluorescence loss kinetics based on a multi-compartment model. However, this method lacked a proper description of intracellular diffusion and nucleo-cytoplasmic exchange of GFP-mtHtt not associated with the IBs [7].

Here, we present what we believe is a new computational method to directly infer the diffusion constants and nuclear membrane permeability coefficients of polyQ proteins as well as their binding dynamics to IBs in concert with bleaching coefficients for the intended laser bleach in the FLIP experiment directly from experimental confocal FLIP images. For that, we made use of a reaction-diffusion multi-compartment model implemented into FEniCS and solved that on a meshed surface geometry directly obtained from the cell images in the FLIP sequence. We used a discontinuous Galerkin (DG) model for improved boundary description and numerical integration of the underlying partial differential equation (PDE) system after transforming that into the weak form.

## Methods

### A reaction-diffusion model on real cell geometry

In [28] we present a reaction-diffusion model with semipermeable nuclear membrane and hindrance for spatial heterogeneity. As described in [28] there is currently put a lot of research effort on understanding the architectures and molecular crowding in living cells. Therefor the computational FLIP model also allows this by a space dependent first order reaction kinetic given by:

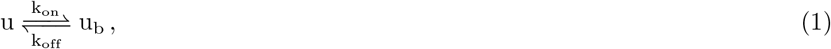

where *u* and *u_b_* is the intensities of the free and hindered molecules, respectively. Letting the observed fluorescence intensity from the FLIP images be described by:

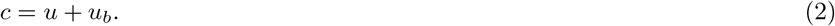

For areas with high intensity we would find a higher population of the hindered *u_b_* proteins. Then given the first order reaction kinetic (1), the space dependent reaction rate *k*_on_ will be high in high-intensity areas and zero in the areas with lowest intensities. Thus letting *c*^0^ be the observed intensity from the first FLIP image, *u*^0^ be the intensity of the free molecules and 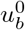 be the intensity of the hindered molecules such that (2) is fulfilled. Letting γ be the proportionality constant then by [28] the reaction rates are set to:

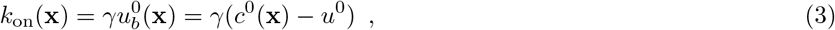

where γ is a proportionality constant. Consequently, *k*_off_ is constant

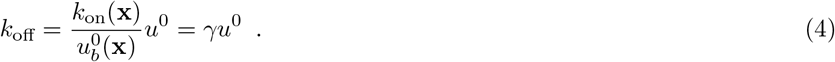

Letting eGFP diffusion be expressed in the terms of Fick’s law and *α* being the diffusion constant for the free eGFP molecules, our time-dependent PDE model reads:

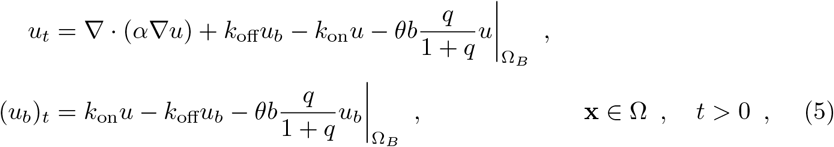

where *θ* is the time dependent indicator function simulating the high intensity laser bleaches, *b* is the intrinsic bleaching rate constant and *q* is the equilibrium constant for the reaction between the ground and excited state for a fluorophore [29]. For mass conservation the Neumann boundary condition along *∂*Ω is used,

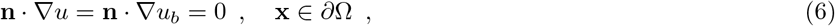

where **n** is the outward unit normal. With initial conditions:

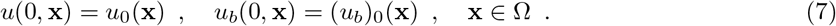

The cytoplasm and nucleus are separated by the nuclear membrane Γ*_M_* with diffusive transport for eGFP through the nuclear pore complex leading to the compartment model presented in [28], where the diffusive flux is expressed as interface condition

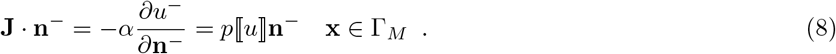

### Multi-compartment modeling of eGFP-mtHtt exchange

As the aggregates cannot always be seen on the first FLIP image, the structure creative reaction mechanism from (1) do not form the aggregates. In [7] a multicompartment model of eGFP-mtHtt exchange between cytoplasm and aggregates where presented. The multi-compartment approach is here transferred into a multicompartment model with a transport process as an internal interface conditions, with the first order transport kinetics described as:

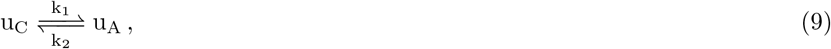

where *u_C_* is the intensity in the cytoplasm and *u_A_* is the intensity in the respective aggregate. Expressed as a differential equation the mass preserving transport process becomes:

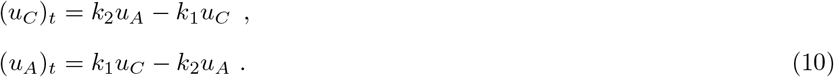

Applied as an interface condition, *u_C_* and *u_A_* becomes the intensities in each of the illustrated neighboring triangles in Figure 1, which are located at the cytoplasmic and aggregate side of the aggregates boundary Γ*_A_*, respectively. Therefore this reaction only happens between two adjacent triangles where their common edge is a part of the line that separates the cytoplasm and aggregates.

**Figure 1.**
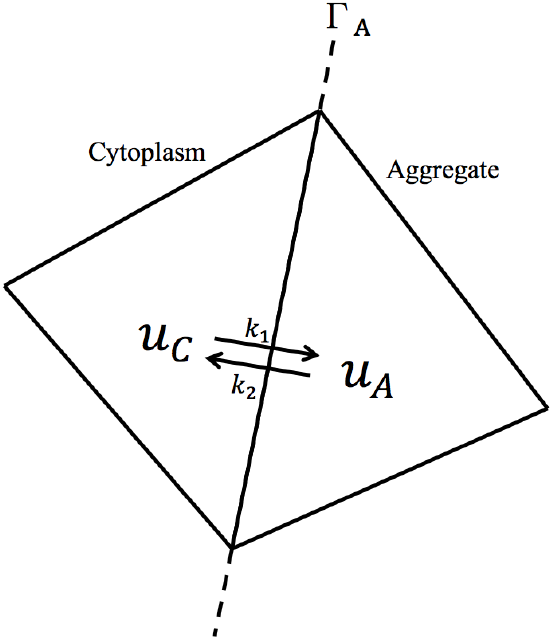
Transport kinetics between the aggregates and cytoplasm.

### Real cell geometry

The cell geometry (see Figure 2) is conveyed from the FLIP images by use of an extended implementation of [30] which uses the “Active Contours Without Edges” method by Chan and Vese[31]. The Chan-Vese model does not depend on the image gradients, and is therefore able to accomplish a segmentation on more blurred images. This Chan-Vese model uses the level set function to iteratively minimize the Chan-Vese energy function that considers the length of the contour and the divergence in the pixel values inside and outside the contour, respectively. As bleaching of the FLIP images occurs in the nucleus, it is hard to segment it automatically from the FLIP sequence. Thus the geometry of the nucleus is here set by hand. However, the cell geometry is segmented from the first image and the aggregates are all segmented from the last FLIP image. The mesh is generated on the geometry in Figure 2 with Gmsh and then converted to XML-file.

**Figure 2.**
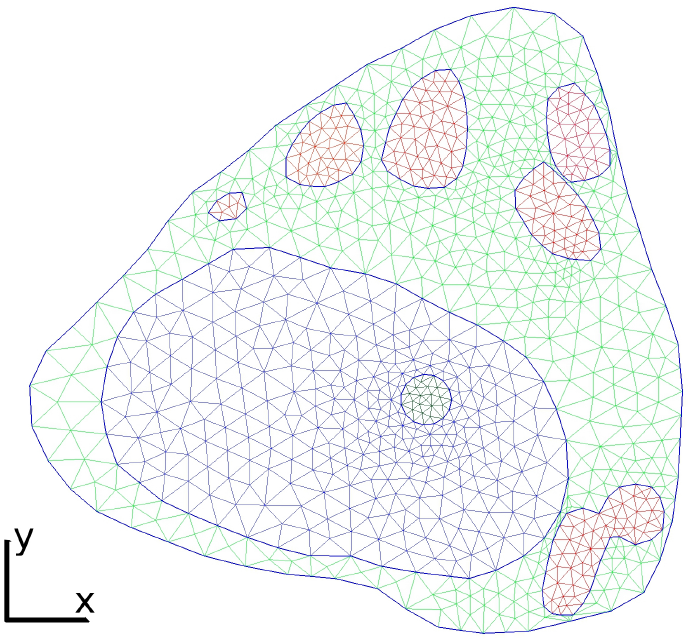
Mesh with 1825 triangles on the real cell geometry. The green triangles constitute the cytoplasm, in red is the aggregates, the dark blue triangles form the nucleus and inside nucleus the round bleaching area with a diameter of 25 μm can be found.

### A discontinuous Galerkin method with internal interface condition

In [28], the interface condition along the nuclear membrane (8) was implemented into the IPDG method based on [32, 33]. Additionally, in this paper, the internal interface condition along the aggregates boundaries are implemented. For the implementation, the weak form for the aggregate interface conditions is here considered.

First let the discretization of Ω be denoted by *𝒯_h_* consisting of disjoint open elements 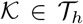. While integrating along Γ*_A_*, *u*^−^ and *u*^+^ are considered as the values of two different but adjacent elements 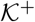 and 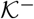 with a common edge on Γ_A_. To rewrite (10) into integral form with the *u*^−^ and *u*^+^ notation, (10) is split up in two cases, one if *u*^−^ is in the cytoplasm and one if *u*^−^ is in the aggregate. An indicator function *I_C_* is therefore introduced as:

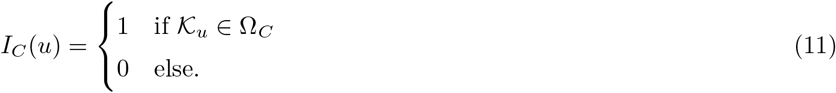

Thus the weak form reads:

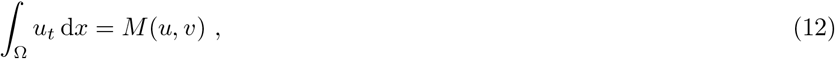

Where

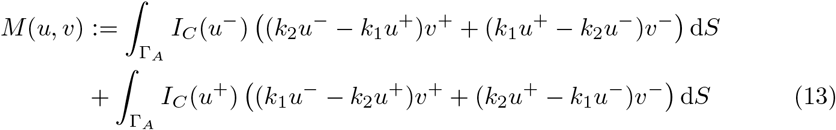

and *v* as the usual test function.

For notation, now let Γ denote the union of the boundaries of all the disjoint open elements 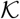. Furthermore, let Γ consist of four disjoint subsets, such that Γ = ∂Ω∪Γ_int_∪Γ*_M_*∪Γ*_A_*. Thus Γ_int_ holds all internal edges. Then usual average and jump term for DG-methods are defined as {*u*} = (*u*^+^ + *u*^−^)/2, ⟦*u*⟧ = *u*^+^**n**^+^ + *u*^−^**n**^−^. For vector valued functions **q** the average and jump term are defined as: {**q**} = (**q**^+^ + **q**^−^)/2, ⟦**q**⟧ = **q**^+^ · **n**^+^ + **q**^−^ · **n**^−^. where **n**^±^ is the outward unit vectors on 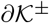.

Reusing the notation from [28] we let

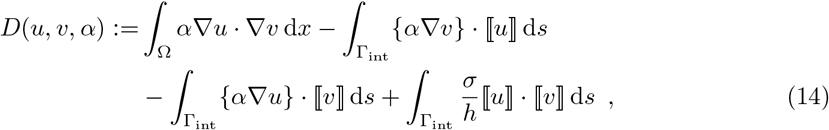

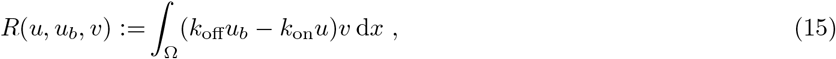

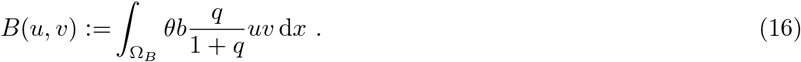

Thus our weak formulation reads:

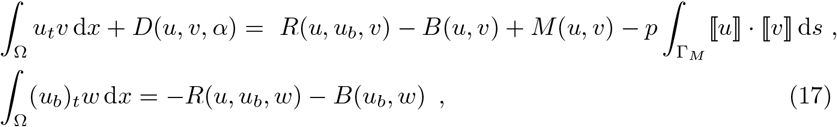

where *v* and *w* are the usual test functions.

Any L-stable method can be used for discretizing the time derivative. Here the backward Euler is used for the implementation using the automated Finite Element package FEniCS [34]. Pre-assemble the system matrix will improve the computational time in FEniCS. However, as the bleaching term is time dependent the system is here pre-assembled into two system matrices. One with and one without the bleaching term. Inside the python script, the weak formulation is therefore expressed twice in the UFL form language.

For simplicity the bleaching term 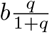 from (16) is replaced by *β* in the implementation and calibration. An example of the weak formulation with the bleaching term is presented here:

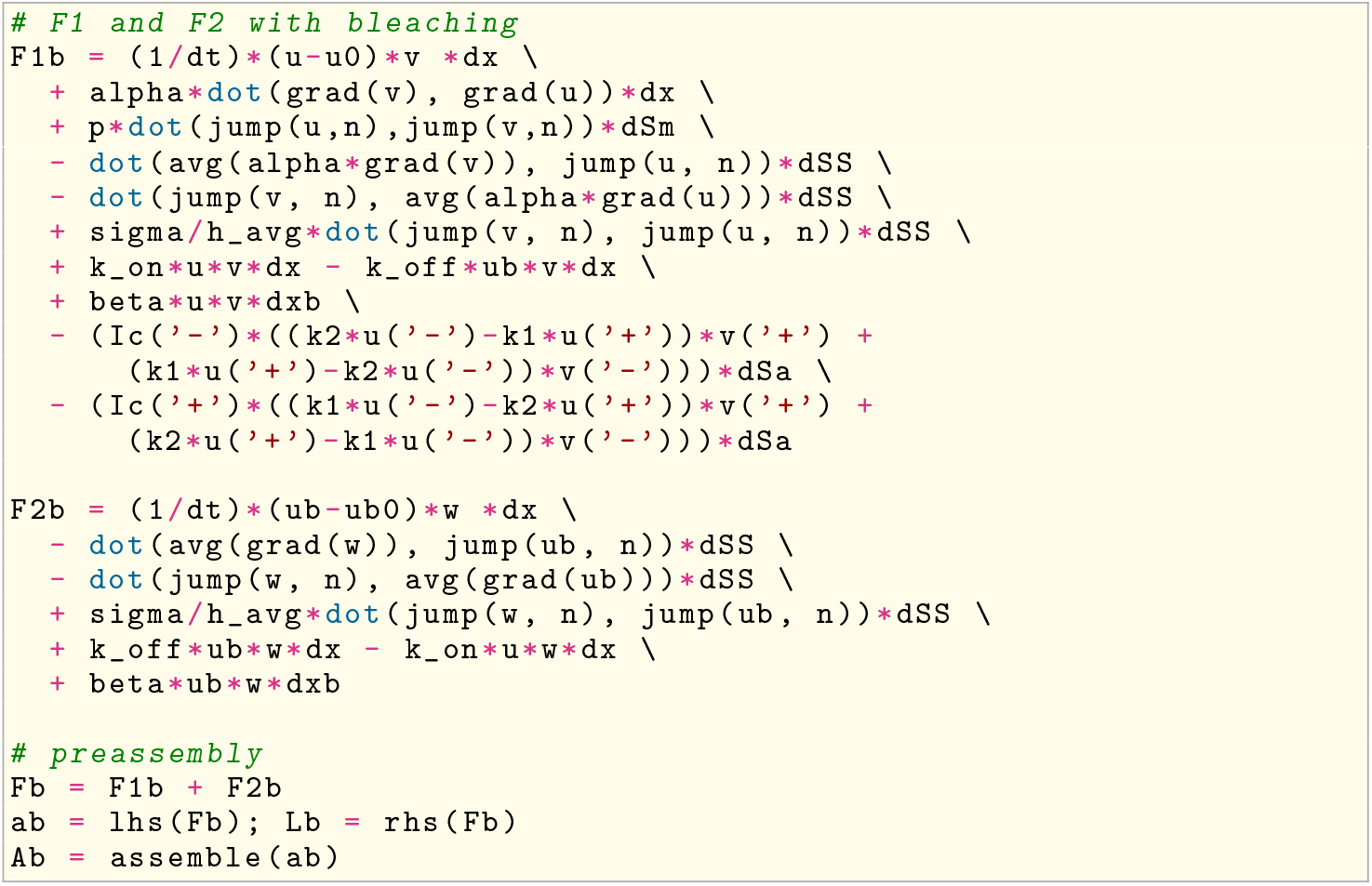

Where `dSm` represent the integral along the membrane, `dSa` is the integral along the aggregates boundaries, `dSS` is the integral on the remaining edges with smooth solutions and `dxb` represents the bleaching area. A similar system matrix is implemented without the bleaching term and the left-hand side is pre-assembled as the matrix A with the right-hand side L. The time dependent system is solved in FEniCS by:

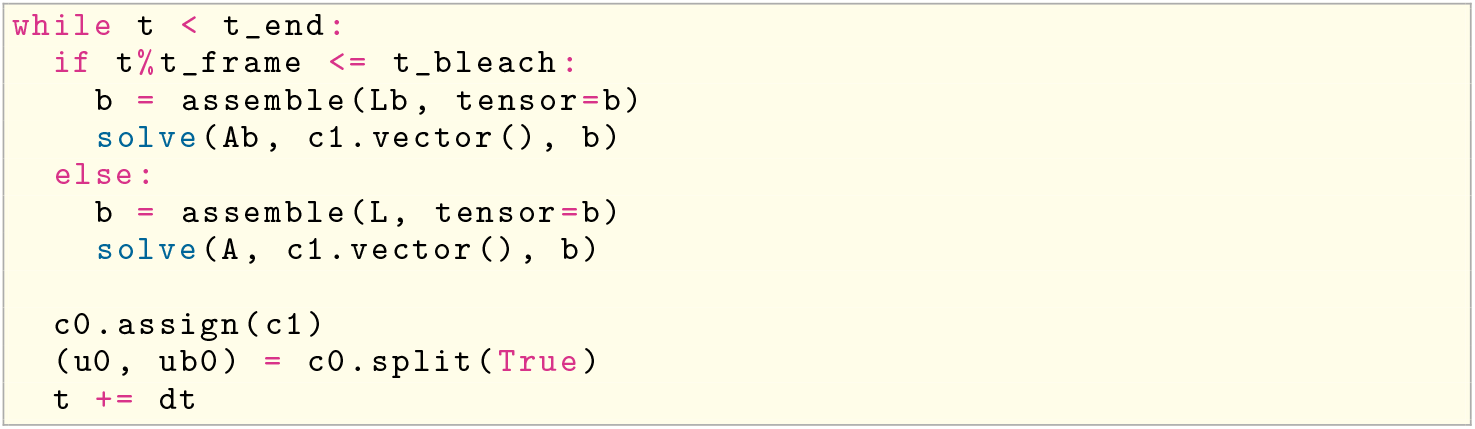

## Results

### Calibration and simulation of intracellular transport

To calibrate the unknown parameters *α*, *β*, *γ*, *p*, *k*_1_, *k*_2_ we make a comparison between the simulation result and the FLIP images. The frame time for the FLIP experiment in Figure 3(A-D) where Δ*t_frame_* = 2.8s, within that time the bleaching area with a diameter of 25μm where bleached with 100% laser intensity for 2s. Thus the imaging process with a laser power of 0.5% took 0.8s.

**Figure 3.**
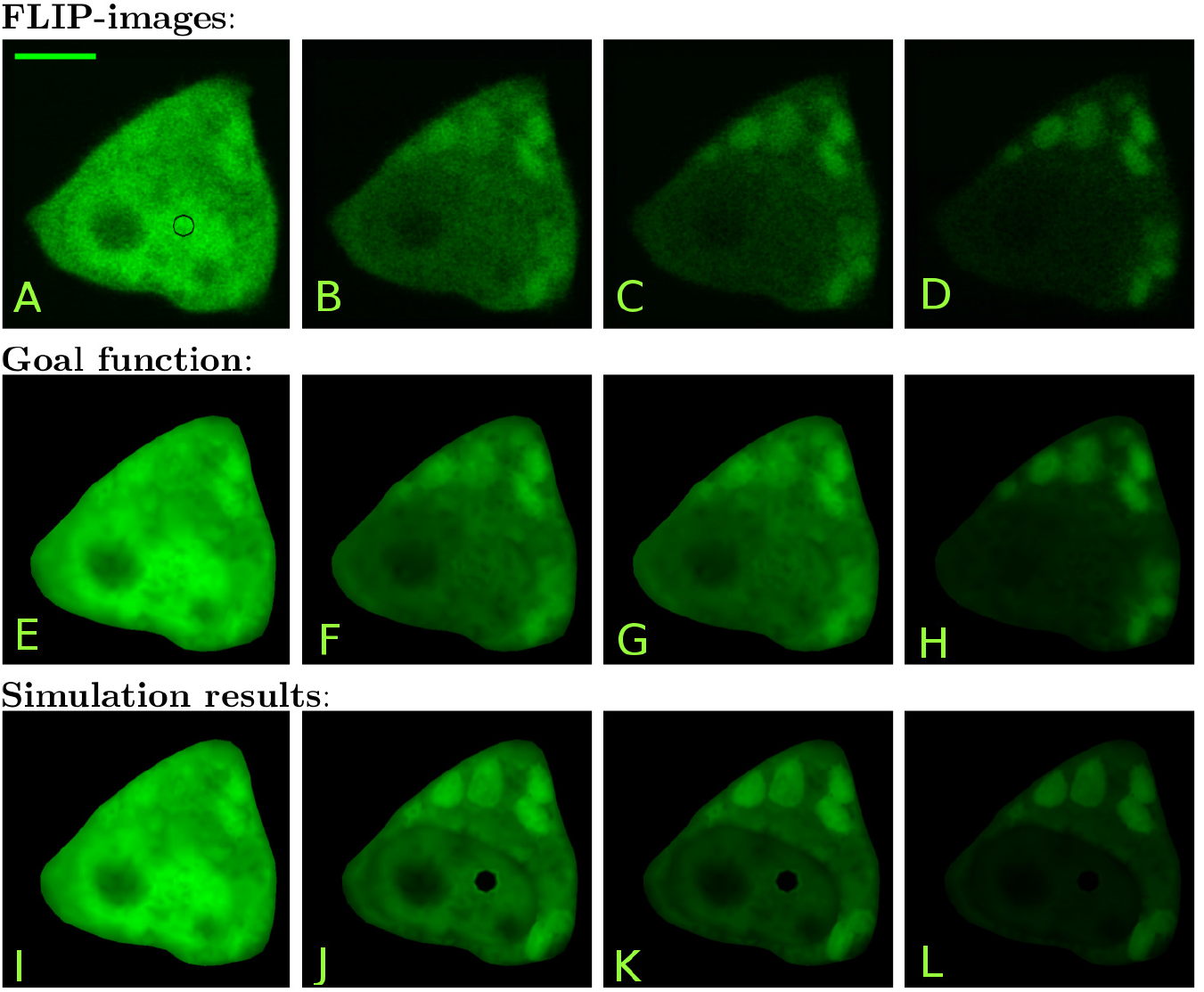
The first four images (A-D) are the original FLIP images of the CHO cells expressing GFP-Q73 in the cytoplasm and nucleus. It is produced in a temperature controlled (35 ± 1°C) environment on a Zeiss LSM 510 confocal microscope using the 488nm line of an Argon laser. The black circle on the image (A) shows the 25-pixel wide bleaching area and a scalebar which is 5 μm. The leftmost FLIP image (A) is taken before bleaching, the next image (B) is taken after it has been bleached 10 times, i.e., time *t* = 28 s. The third FLIP image (C) is the 20’th FLIP image in the sequence (time *t* = 56 s) and the last (D) is at time *t* = 109.2 s which correspond to FLIP fame 39. The second row (E-H) shows the corresponding goal function. The third row (I-L) shows the simulation results, all at times corresponding to the displayed FLIP images.

To easily compare the simulation results and the FLIP sequence, the goal function seen in Figure 3(E-H) is created. The goal function is a piecewise linear discontinuous Galerkin function defined on the mesh, which represents the values from the denoised FLIP images. To denoise the FLIP sequence, Gaussian blur with a radius of 1 pixel is used. At the discrete times *t_i_* = Δ*t_frame_*(*i* – 1) + *t_compare_* seconds *i* = 1, 2, 3,…, *n* the L_2_ norm of the difference between the goal function and the simulation is calculated to represent the misfit functional as:

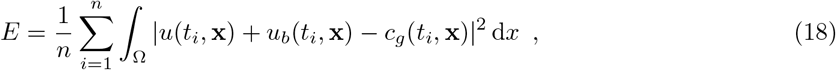

where *c_g_* is the goal function. For the sequence in Figure 3 the number of FLIP images is *n* = 40 and the time where the simulation and FLIP data are compared is *t_compare_* = 2.6s. To calibrate the unknown parameters, the Nelder-Mead downhill simplex algorithm [35] from the SciPy library [36] is used. The stop criterium is set such that either the difference in the parameter or the difference in the misfit functional between each iteration should be lower than 10^−4^. Looking at the reactions rates *k*_1_ and *k*_2_ it is known from (10) that in equilibrium the equilibrium constant can be described as:

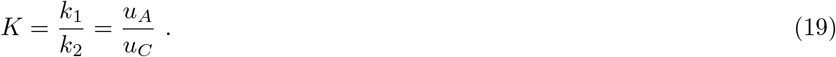

Assuming that the first FLIP image before bleaching (see Figure 3A) is in equilibrium, *K* can be determined by the use of the average intensities from inside the aggregates and cytoplasm, respectively. From the FLIP image in Figure 3A the equilibrium constant turns out to be *K* = 1.16. Thus by expressing *k*_2_ in terms of *k*_1_, the parameters that need to be calibrated are reduced to *α*, *β*, *γ*, *p*, *k*_1_. The initial guesses for the calibration are set to *α*_0_ = 25, *β*_0_ = 20, *γ*_0_ = 0.5, *p*_0_ = 0.05 and (*k*_1_)_0_ = 0.001. After 405 iterations and 679 evaluations, the resulting calibrated parameters are

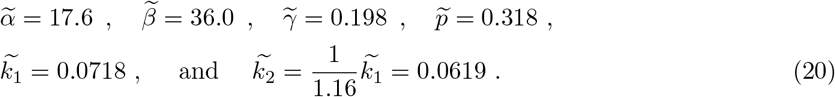

The misfit functional with the initial parameters *E*_0_ = 7,141 where lowered to *E* = 2,807 for the calibrated parameters in (20). The calibration process took around 9 hours on an Intel Core i5 processor at 3.2 GHz with 8 GB memory running Ubuntu 16.04 LTS. The results of the calibration process are presented in Figure 3(I-L).

In Figure 4(A-D) a similar FLIP sequence with Δ*t_frame_* = 2.6s, *t_compare_* = 2.4s and *n* = 55 can be seen. The simulations have been made on a mesh consistent of 1998 triangles, and the initial guesses for the calibration are set to *α*_0_ = 15, *β*_0_ = 10, *γ*0 = 0.05, *p*_0_ = 0.5 and (*k*_1_)_0_ = 0.01. After 179 iterations and 293 evaluations within five and a half hour the resulting calibrated parameters are

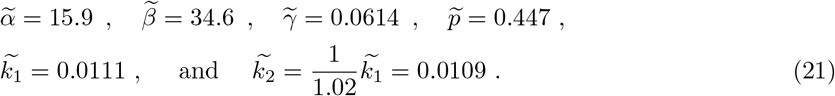

**Figure 4.**
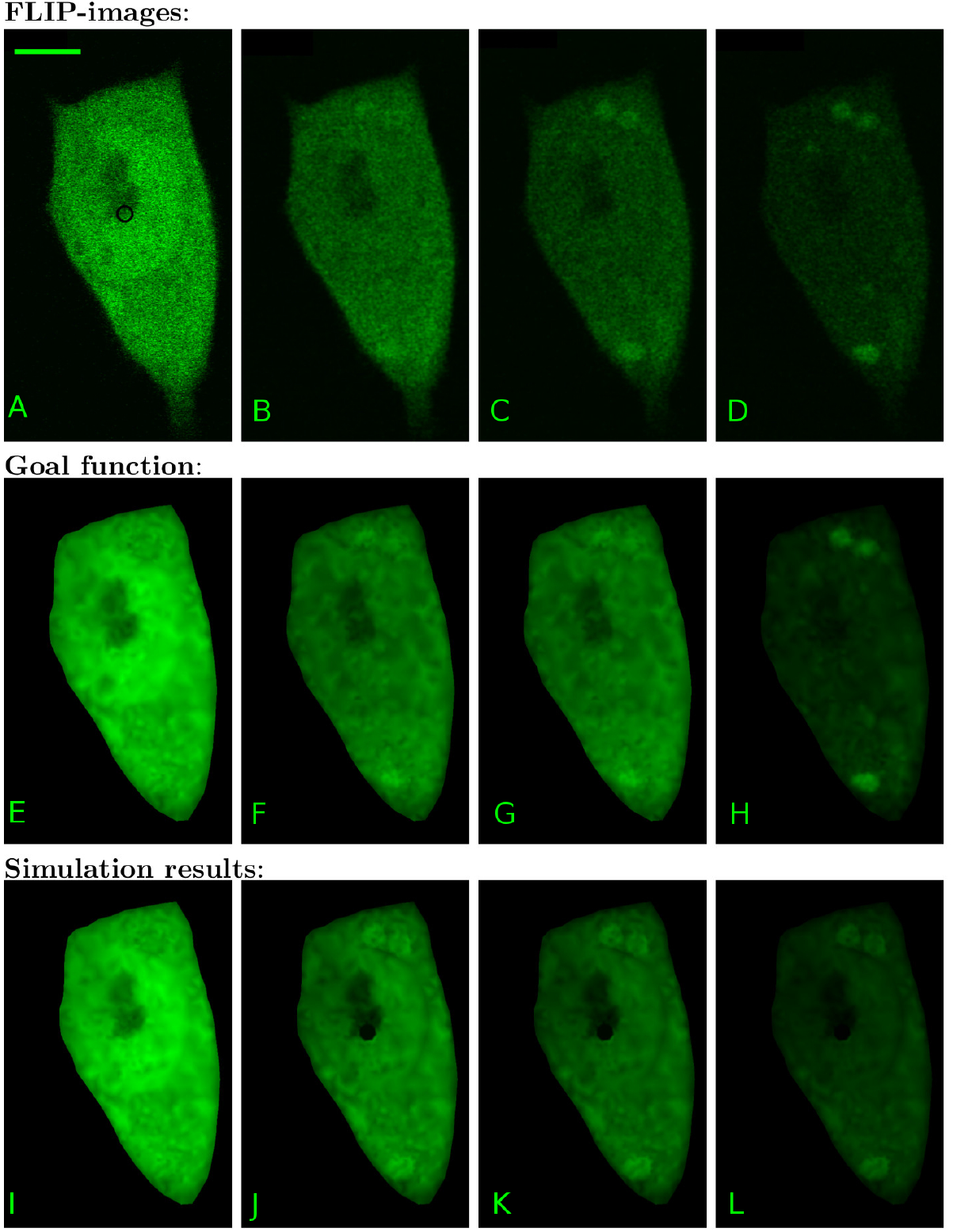
(A-D) are the original FLIP images of the CHO cells expressing GFP-Q73 in the cytoplasm and nucleus. The black circle on the image (A) shows the 18-pixel wide bleaching area and a scalebar which is 5 μm. (A) is taken before bleaching, (B) is after 10 time bleaches, i.e., time *t* = 26 s. (C) is the 20’th FLIP image in the sequence (time *t* = 52 s) and (D) is produced at time *t* = 104 s which correspond to FLIP fame 40. The second row (E-H) shows the corresponding goal function. The third row (I-L) shows the simulation results, all at times corresponding to the displayed FLIP images.

The simulation result with the calibrated parameters can be seen in Figure 4(I-L).

### Calibration test

To test the calibration approach a forward simulation with known parameters is made to represent and replace the FLIP images, which we calibrated against. The forward simulation is made with the same initial and boundary conditions as used in Figure 3, on the mesh from Figure 2. The chosen parameters are:

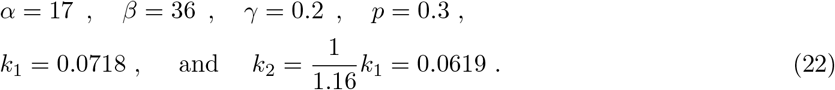

Gaussian noise with the mean set to zero and a variance whose size is approximately 10% of the maximum intensity is added to the results of the forward simulation. The forward simulation result now replace the goal function that is usually extracted from the experimental FLIP images in the calibration process. The rest of the setup, including the initial guesses on the parameters for the calibration, is identical to the one used for the calibration in Figure 3. Through the calibration process the misfit function *E* was lowered from 639.4 to 169.7 in 388 iterations with 612 function evaluations which took around 10 hours. The calibrated parameters are:

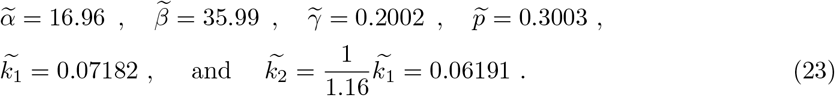

A small error is seen on the fourth digit, which is due to both the Gaussian noise and the size of the stop criterion for the Nelder-Mead algorithm.

## Discussion

Phase separation and aggregation of polyQ proteins are prominent signs of certain neurodegenerative diseases. Often, protein inclusions of GFP-tagged polyQ proteins are first visible in cells after several days in culture allowing only for studying relatively inert, bright and stable aggregate structures [15, 13]. Thus a key requirement in traditional approaches is that the IBs and similar fluorescent protein aggregates differ in their intensity significantly from the fluorescent protein pool in the surrounding cyto- or nucleoplasm. This, however, limits the analysis to certain inclusion types. Here, we present a new computational approach for inferring diffusion, membrane permeability, and exchange rate constants of GFP-mtHtt between cytoplasm and aggregates of differing brightness directly from experimental FLIP image sequences. Our method allows for detection and dynamic characterization of protein aggregates even in cases, where they are not visible in single image acquisitions. Using the calibrated reaction–diffusion model, we found that rate constants for exchange of GFP–mtHtt between such large but dim inclusions and the cytoplasm are fast (binding rate constant *k*_1_ = 0.0718 s^−1^ (Figure 3) and *k*_1_ = 0.0111 s^−1^ (Figure 4) and release rate constant of *k*_2_ = 0.0619 s^−1^ (Figure 3) and *k*_2_ = 0.0109 s^−1^ (Figure 4). We found similar values previously for the same protein and cell system using a simple multi-compartment model which ignored diffusion and nucleo-cytoplasmic exchange of GFP-mtHtt (i.e. binding rate constant *k*_1_ = 0.016 ± 0.006 s^−1^ and release rate constant of *k*_2_ = 0.0127 ± 0.004 s^−1^, mean ± SEM of 6 cells) [7]. From that, we can conclude, that the typical residence time of GFP-mtHtt once bound to cytoplasmic aggregates is on order 16-83 s before being again released and available for free cytoplasmic transport and nucleo-cytoplasmic exchange. Our estimates of intracellular diffusion constants for GFP-mtHtt of 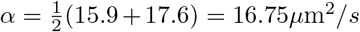 are in good agreement with what would be expected for a protein the size of GFP-Q73 (i.e. Stokes radius of R ≈ 3.4 nm [9]) in the cytoplasm (i.e. viscosity of 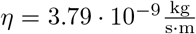 predicts α = 16.6 according to data from [37]). Supporting that notion is a previous report, which found *α* = 18.4 ± 3.3*μ*m^2^/*s* for diffusion of GFP-mtHtt of the same size (i.e., Q73) in the cytoplasm of N2a cells using FRAP [9]. Using an average cytoplasmic diffusion constant of *α* = 16.75*μ*m^2^/*s* and the upper estimate of the time constant for binding of 1/(*k*_1_ = 0.0718 s^−1^) = 14 s from our analysis, we conclude that GFP-mtHtt can diffuse on average 30*μ*m away from an aggregate after release before the next binding event takes place. Thus, diffusion is not limiting the aggregation kinetics, which explains, why we found very similar estimates for the binding and dissociation constants as reported here with our previous model which ignored cytoplasmic diffusion altogether [7]. We believe that rapid diffusion and exchange of soluble mtHtt with cytoplasmic inclusions could contribute to the efficient recruitment of other proteins to IBs which further accelerates cellular dysfunction as observed in various studies [38, 14, 6].

In [28] we presented a method using a semi-permeable membrane model to describe the transport of eGFP. The same semi-permeable membrane model is used in this paper. However, the relatively high permeabilities of the GFP-mtHtt protein may indicate that the traffic across the nuclear membrane could be caused by selective and directed transport [39]. In fact, we found that nuclear membrane permeabilities for GFP-mtHtt in the cells studied in Figure 3 and 4 were higher than what we previously observed for GFP using the same FLIP modeling approach (ref. Scientific Reports-MS). On the other hand, two to three days after transient transfection, we often observed slowed nuclear-cytoplasmic exchange of GFP-mtHtt compared GFP, likely due to the pronounced formation of sub-resolution aggregates which interfere with normal nucleo-cytoplasmic transport (not shown but see Figure 6 in [7]). Such varying results have been reported previously [40, 27, 41, 42, 43] and they could be well attributed to the eventual occurrence of soluble oligomers, whose transport across the nuclear membrane is delayed, while transport of monomeric mtHtt profits from interaction with FG-rich repeats in the nuclear pore, which can accelerate transport compared to passive cargo [37]. It is possible to replace the semi-permeable membrane model with a model similar to the compartment model for the aggregates, and thus obtain a reactive membrane transition.

The proportionality constant *γ* depends on the structures and intensity observed on the first FLIP image. Those these cannot be expected to be fully equal, as they are structure dependent.

In this paper, only one reaction rate is fitted for all aggregates in the same cell. Each extra reaction rate per aggregate would increase the complexity of the calibration process, such that one should have independent evidence for such heterogeneity before extending the model into that direction. For the readers that may want individual reaction mechanics for each aggregate, we suggest to calibrate the parameters *α*, *β*, *γ* and *p* first, and then fix these parameters while finding the ones for the aggregates. This can be done under the assumption that the traffic from the aggregates is so small that it would not affect the other parameters.

## Conclusion

Our new computational method allows one to determine diffusion constants, nucleo-cytoplasmic permeability and exchange kinetics of polyQ proteins, such as mtHtt, from live-cell FLIP image data. This is the first time, to our knowledge, that all such transport parameters can be inferred in parallel from the full spatiotemporal FLIP intensity profile directly within the cell geometry. Using this new method, we find that polyQ proteins can exchange rapidly between cytoplasm and aggregates and that diffusion of protein monomers is not limiting this exchange process. Furthermore, we show that computational FLIP is an efficient method to detect dim protein aggregates due to their delayed fluorescence loss. Binding and dissociation constants of mtHtt to and from such aggregates are comparable such that the inclusions are hardly visible in single images. Finally, our method sets the stage for a systematic exploration of how the aggregation process affects the nucleo-cytoplasmic permeability of polyQ proteins. Our new approach is widely applicable to quantify protein dynamics in cellular inclusions of various disease models.

